# The costs of close contacts: Visualizing the energy landscape of cell contacts at the nanoscale

**DOI:** 10.1101/697672

**Authors:** K. Kulenkampff, A. H. Lippert, J. McColl, A. M. Santos, A. Ponjavic, S. F. Lee, S. J. Davis, D. Klenerman

## Abstract

Cell-cell contact often underpins signalling between cells. Contact is mediated by proteins on both cells creating interfaces with gap sizes typically around 14 nm. Protein binding and accumulation leads to the contact becoming crowded, reducing the rate of protein diffusion, even for unbound proteins. Here we show that, by tracking quantum dots of different dimensions for extended periods of time, it is possible to obtain the probability of a molecule entering the contact, the change in its diffusion upon entry and the impact of spatial heterogeneity of adhesion protein density in the contact. By analysing the contacts formed by a T cell interacting with adhesion proteins anchored to a supported lipid bilayer, we find that probes are excluded from contact entry in a size-dependent manner for gap-to-probe differences of 4.1 nm. We also observe probes being trapped inside the contact and a decrease in diffusion of up to 85% in dense adhesion protein contacts. This approach provides new insights into the nature of cell-cell contacts, revealing that cell contacts are highly heterogeneous, due to topography- and protein density-related processes. These effects are likely to profoundly influence signalling between cells.

**Statement of Significance:** The spatial distribution and diffusion of proteins has been shown to be important for various signalling machineries. As such size-dependent reorganisation of proteins in the immune cell-contact has been shown to affect activation of immune cells. While these studies relied on bulk measurements to investigate protein exclusion, small scale topographical changes and protein dynamics could not be evaluated. However, recent studies show that T cell activation is mediated by nanoscale structures. In our study the use molecular probes of various sizes to investigate the energy landscape of single molecules in a cell contact. This provides additional information and insights which cannot be determined by performing bulk experiments alone indicates.

## Introduction

Cells in multicellular organisms are continually in contact with other cells and signal through a variety of mechanisms (1–3). Juxtacrine or cell-cell signalling is a process wherein two cells form a sustained contact allowing molecules on the cell surfaces to interact and exchange at the cell-cell contact. In the case of lymphocytes, fragments of pathogens presented by antigen presenting cells (APCs) engage the T-cell receptor (TCR) across these contacts, leading to signalling and T-cell activation, and eventually an immune response. Activation results in the large-scale spatial reorganisation of other important membrane proteins, including signalling and adhesion proteins, into a structure called the immunological synapse (4).

Although we understand very little about the mechanisms which lead to this restructuring, the dimensions and steric properties of surface proteins are thought to be important in their spatial arrangement (5, 6). In this context, it has been suggested that passive rearrangements alone could explain TCR triggering (7, 8) (9, 10). The contact gap between the two cells is set by adhesion molecules, matching the dimensions of the TCR/ligand complex at ~14 nm. The extracellular domains of other signalling proteins, such as the inhibitory phosphatase CD45, extends well beyond this distance. In the contact, CD45 would experience size-dependent exclusion and the lack of inhibition at the contact is proposed to contribute to signalling (7, 11, 12). In addition, protein-protein interactions (13), putative lipid rafts (14) as well as the cytoskeleton (15, 16) have been shown to influence the diffusional behaviour of proteins, with effects on signalling (13). Finally, the diffusional behaviour can also be changed by steric confinement imposed by contact formation and hence proteins inside the contact will experience crowding and physical restrictions.

To understand the relevance of steric exclusion and the contribution of physical restrictions on protein diffusion in contacts, it is important to develop methods for analysing how the contact gap size affects protein behaviour at the single molecule level. Here, using a probe of known dimensions, we have been able to study the effect of steric hindrance and crowding experienced by proteins at the contact. Previous studies used small fluorescent molecules, including sugars (17), fluorescent proteins (18) and Quantum Dots (QDots) (19, 20) to probe cell-cell, cell-surface or membrane-membrane contacts in bulk experiments. They identified a size-threshold leading to partial-to-complete clearing of the immune synapse or contact for particles 2-10 nm larger than the expected contact. While these experiments yielded important insights into the general size-excluding properties of contacts, it has remained unclear how contact topography, particle exchange across contact boundaries, and energy penalties for entering and exiting the contact influence protein organization at cell contacts.

## Materials & Methods

### Cell Culture

The Jurkat rCD48 cell line was generated via lentivirus transfections. T cells were cultured in phenol red– free RPMI supplemented with 10% FCS, 1% HEPES buffer, 1% sodium pyruvate and 1% penicillin-streptomycin.

### Supported Lipid Bilayer (SLB) Formation

SLBs were prepared according to previous protocols (21). Glass cover slides (VWR, International) were cleaned for one hour using piranha solution (3:1 sulfuric acid/ hydrogen peroxide). After rinsing with MiliQ water the slides were plasma cleaned for 30 minutes and a silicon well (Grace Bio-Labs) attached to each slide. Previously prepared small unilamellar vesicle (SUV) solution was added to each Phosphate buffer saline (PBS) filled silicone well and incubated for 30 minutes. The vesicle solution consisted of 1 mg/ml of 95% POPC and 4.999% DGS-NTA(Ni) and 0.001% Biotinyl-Cap-PE (all Avanti Polar Lipids, Alabaster USA). After incubating for 30 minutes the wells were washed three times with PBS solution and the protein solution added at 30 μg/ml. Purified protein spacers were gratefully provided by the Davis lab in Oxford. The proteins used were either rCD2.D1, a truncated protein with domain 1 of rCD2 or a chimeric protein that comprises the extracellular portion of rCD2 plus rCD45 (rCD2rCD45). Each of the proteins was labelled with Alexa Fluor 647 (Thermofisher). All experiments carried out at nickel lipid saturation and all bilayer conditions showed similar protein fluorescence intensities. Variations of maximal 11% s.d. within one condition and mean variations of maximal 15% s.d. between the conditions. (see Suppl. Figure 1). After 1hr incubation at room temperature the wells were washed three times with PBS and cells added.

### Sample Preparation

Streptavidin coated QDots (QDot 525 Streptavidin Conjugate, QDot 605 Streptavidin Conjugate, both Thermo Fisher Scientific) were suspended onto the bilayer at a concentration of 50 nM in PBS. The QDots were left for 5 min before washing two times with PBS to remove QDots that had not bound to biotinylated headgroups.

### Data Acquisition

Prior to imaging the cells were washed in PBS and then added to the rCD2 constructs and QDot containing bilayer (preparation described above). Imaging was performed using a custom-built TIRF setup using a 100 × Apo TIRF, NA 1.49 objective, (Nikon, Japan) creating a TIRF illumination at the glass water interface. Fluorescence was recorded through a beam splitting system (Dual-View, BioVisionTechnologies, US) using a dichroic mirror and filters (for 488/633 emission FF605-Di02 (dichroic, Photometrics, US), FF03-525/50-25 (filter, 488 emission) and BLP01-635R-25 (filter, 633 emission), all Semrock, US). 1000 frames per experiment were acquired with an exposure time of 30 ms. The camera (Cascade II,Photometrics, US) and shutter (SH05, Thorlabs, US) were operated using Micromanager (Vale Lab, US). The two DualView channels were aligned using TetraSpec Microspheres (0.1μm, fluorescent blue/green/orange/dark red, ThermoFisher, US) routinely to a precision of ~120nm.

### Data Analysis: Single-Particle Tracking

Single-Particle Tracking was performed using a custom-written Matlab code (22), yielding diffusion behaviour and coefficients from JD distributions of the QDots. Here, only tracks longer than 5 frames were used with mean track lengths of 14.3±2.3 and SNR of 8.0±1.1.

### Exclusion Analysis

To analyse the size-dependent exclusion, contact masks were defined by thresholding the gradient of the fluorescence of protein spacers (threshold set to 0.2 times the maximum gradient of an image) using a custom written Matlab code. Furthermore, a small size threshold was applied to exclude bilayer inhomogeneities. These masks were used to distinguish between QDots inside and outside the contact by performing peak find on the first frame of every experiment to calculate the ratio of density inside vs outside the contact. (Fiji: an open-source platform for biological-image analysis).

### Energy Penalty

We used the contact masks to divide tracks and jumps into four categories: 1, tracks that are outside the contact mask, which stay outside the contact mask. 2, tracks that are outside the contact mask and end inside the contact mask. 3, tracks, which start inside the contact mask and stay inside or 4, tracks, which start inside the contact mask and end outside the contact mask. To avoid bias towards unsucessfull entries or exits of QDots, we analysed only tracks which were recorded for at least five frames after touching the cell border. Hence, we could analyse parameters, such as jump distance in the corresponding location category, number of attempts of a QDot to enter cell gap and number of successful entries into or exits out of the gap.

Using those success rates, we could calculate energy penalties, *ε*_*i*_, for entering and exiting the contact via the Boltzmann distribution, 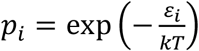, with 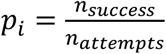 to calculate *ε*_*enter*_ or *ε*_*exit*_

### Size- and density dependent slow down

Using the track ensemble, we created a map of the average speed per pixel of diffusing QDots. The track ensemble was overlaid with a pixel grid corresponding to the original pixel size of the recorded image to determine average jump distance at the corresponding pixel position. These were displayed with scaled colours corresponding to the magnitude of the average jump distance. By fitting the JD distribution jumps occurring inside or outside the contact, we could determine the diffusion coefficient inside and outside the contact, and across different intensity zones (corresponding to different protein densities). JD distribution were fitted to the probability distribution 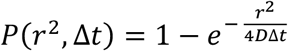 of a particle with a diffusion coefficient, ***D***, to be within the radius ***r*** of a shell at a time **∆*****t*** (22).

### Statistical testing

One-way ANOVA and post hoc tests were used as noted (Prism 8, Graphpad) using a significance level of α < 0.05 for ANOVA and for post hoc tests. The Tukey-Kramer method was applied to evaluate p-values in post hoc tests.

## Results and Discussion

### QDots of various sizes experience size-dependent exclusion

Exclusion of molecules from cell-cell contacts has been proposed as a key factor during lymphocyte signalling (7, 8) and while there is evidence accumulating to support this idea (23–25) it remains contentious. We sought to determine if large, surface anchored molecules (represented by QDots) would be excluded from contacts by studying the size-dependent exclusion of the QDot probes.

We used supported lipid bilayers (SLB) presenting ‘spacer’ adhesion proteins of various sizes, and QDots of known diameter, freely diffusing in the bilayer as probes (see Suppl. Video 1, Suppl. Fig. 2). To control QDot density, biotinylated lipids were incorporated into the SLBs and the spacer proteins were coupled via nickelated lipids. We controlled cell-bilayer gap sizes using modified versions of the adhesion protein rat CD2 as spacer protein (Fig. 1A). CD2 has two immunoglobulin superfamily domains in its extracellular region, stacked on top of one another, with the membrane distal domain binding rat CD48, which we expressed at the surface of Jurkat T cells. The distance spanned by this complex is approximately 13.4 nm (11). To create small gaps we used a short, truncated version of CD2 comprised of only the ligand binding domain (CD2d1). For larger gaps we used the extracellular region of rat CD2 and the folded part of the extracellular domain of human CD45 (CD2CD45), a large receptor-type phosphatase (Fig. 1A). The CD2d1/CD48 complex forms a gap predicted to be 9.4 nm and CD2CD45:CD48 a 34.4 nm gap (11) (26). To avoid any direct signalling effects through CD48 binding, the cells expressed a truncated form of the receptor lacking a cytoplasmic region. The cells were then dropped onto a SLB containing QDot and CD2 variants, allowing formation of ‘spacer’ complexes. While the bound QDots diffuse freely with a diffusion coefficient of around 0.6 µm^2^s^−1^ (see Suppl. Fig 2, Suppl. Video 1), the cells accumulated CD2 and formed contacts. To probe the effect of steric hindrance on probes of known height we used QDots of two sizes, Q605 and Q525, which have hydrodynamic diameters of 21.2 nm (19) and 14 nm (Thermofisher, Germany, personal communication, March, 2019), respectively (Fig. 1B). We termed the difference between the gap created by the spacer complex (P) and the QDot diameter (Q), Δ(P,Q)=P-Q. This value ranged from 11.3 to 14.9 nm, and we expected to observe size-dependent exclusion for negative Δ(P,Q) values (Fig. 1B).

**Figure 1:**
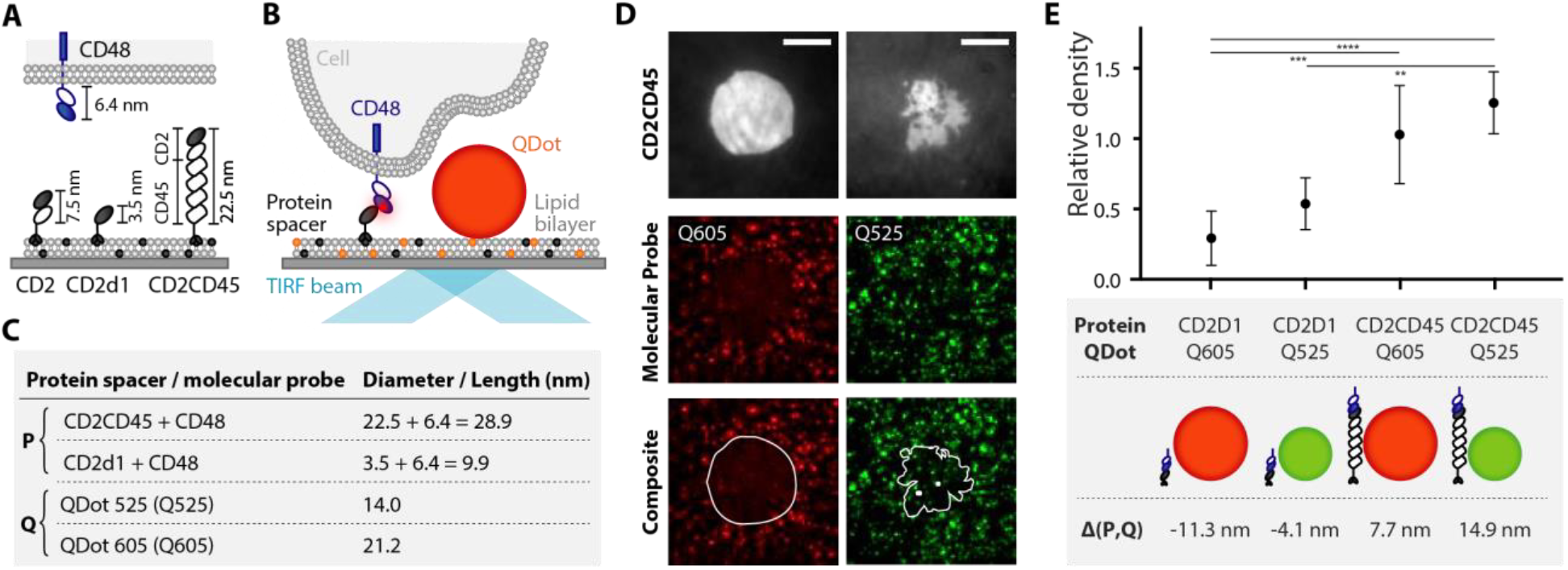
Gap size dependent QDot exclusion. (A) Schematic representation of the protein spacers and their ligand (CD48: 6.4nm) used in the experiments (CD2d1: 3.5nm or CD2CD45: 22.5nm) and the experimental configuration (B): Streptavidin conjugated QDots (Q525 or Q605) were seeded onto a supported lipid bilayer (SLB) containing biotinylated headgroups (orange). Cells form a contact with the SLB by binding protein spacers (P), which is anchored to the SLB through histidine tags on the protein binding nickelated lipids (darkgrey). Spacers, fluorescently labelled with Alexa647, are binding CD48 on the cell, creating gaps of different sizes. (C) The length of each protein spacer and hydrodynamic diameter of QDots used in SLB experiments in this work is presented in the table. (D) TIRF images and composites of the contact formed by CD2CD45 (top) and QDots (middle) in the SLB (right: QDot605 (red), left: QDot525 (green)). Scale bar, 5µm. (E) QDot density inside the cell-bilayer contact are compared to the density outside the contact to gain relative density. Δ(P,Q)=P-Q is the difference between QDot diameter and spacer-protein induced gap size. Number of cells n for each condition: Δ(P,Q)=−11.3 nm, n=6, Δ(P,Q)=−4.1 nm, n=3, Δ(P,Q)=7.7 nm, n=6, Δ(P,Q) =14.9 nm, n=6. Data was analysed using a one-way analysis of variance (ANOVA). **p < 0.01, ***p < 0.001, ****p < 0.0001.

To examine the extent of size-dependent exclusion, we determined the QDot density inside relative to outside the contact (Fig. 1C,D). We observed a lower QDot density beneath the contact region, relative to the density of QDots in the surrounding area, even when the height difference was as small as −4.1 nm (Fig. 1E). The relative density was 29.3±17.5% for the large and 53.7±15.1% for the small QDots in contacts with the smallest gap, giving rise to size-dependent exclusion (exclusion = 1-rel. density) of 46.3% (Δ(P,Q)=−4.1nm) and 70.7% (Δ(P,Q)=−11.3nm). Positive Δ(P,Q) values did not result in observable exclusion effects.

Bulk fluorescence imaging of the distribution of large and small molecules at cell contacts can convey the impression that it is an ‘all or nothing’ effect: proteins larger than the gap are wholly excluded from contacts, whereas proteins smaller than the gap can gain access, but single molecule measurements suggest otherwise (25, 27). Our single-molecule data confirmed that even molecules twice the size of the gap were not wholly excluded (Fig. 1E). We could also observe exclusion effects in contacts 4.1 nm smaller than the QDot (Fig. 1E), which is in good agreement with bulk size exclusion measurements in cell-cell contacts (28) and model membrane systems (18). A difference of 3.2 nm was enough to exclude QDots in other studies (19). This suggests that at cell contacts, size exclusion very likely fine-tunes the distribution of proteins, rather than purely having an “all or nothing” size-exclusion effect.

### QDots are size-dependently restricted from contact entry

Having established that QDots are gap-size dependently excluded, we next studied the entry and exit of QDots from the contact. By tracking single QDots at the border of the contact, we classify the trajectories as entering, exiting and deflected at the contact border (Fig. 2A-C). Tracks starting outside the contact and ending inside it were classified as ‘entering’; tracks exhibiting the opposite behaviour were taken to be ‘exiting’. ‘Deflected’ tracks were classified as tracks starting outside, having one or more localisations on the contact border and ending outside the contact. Using the rate of entries/exists to attempts, we calculated the energy penalty for entering (ε_enter_) and exiting (ε_exit_), assuming a Boltzmann distribution (see Materials & Methods, Fig 2D). For QDots exiting the contact region, ε_exit_ is comparable for all Δ(P,Q) (Fig 2E). Conversely, for QDots entering the contact, ε_enter_ is Δ(P,Q) dependent, with increasing ε_enter_ for decreasing gap-to-probe sizes (Fig. 2E). We observe a significantly larger ε_enter_ than ε_exit_ for negative Δ(P,Q), and unexpectedly also for Δ(P,Q) = 7.7 nm. To confirm this result, we simulated particles with starting positions inside the contact, using contacts from the acquired data set. We allowed free Brownian diffusion, with density and diffusion coefficients adjusted to the experimental data and allowed particles to diffuse freely over the contact. Here, diffusing particles enter and exit the contact without any restrictions, which is shown by the low energy penalties compared to the experimental data. While the simulations confirm that contact entry is restricted, even for positive Δ(P,Q), these values also suggest that QDots exiting the contact also experience a restriction, which is gap-size independent. The spread of ε_exit_ for the experimental data could be an effect of varying protein crowding, which might inhibit exit of proteins from the contact. In addition, by comparing the instantaneous velocity, i.e. jump-distance (JD) distribution of QDots which failed to enter with that of QDots successfully entering the contact, we observed a shift towards lower values for contact gaps with negative Δ(P,Q), also suggesting that there is an energy penalty for contact entry (Suppl. Fig. 3).

**Figure 2.**
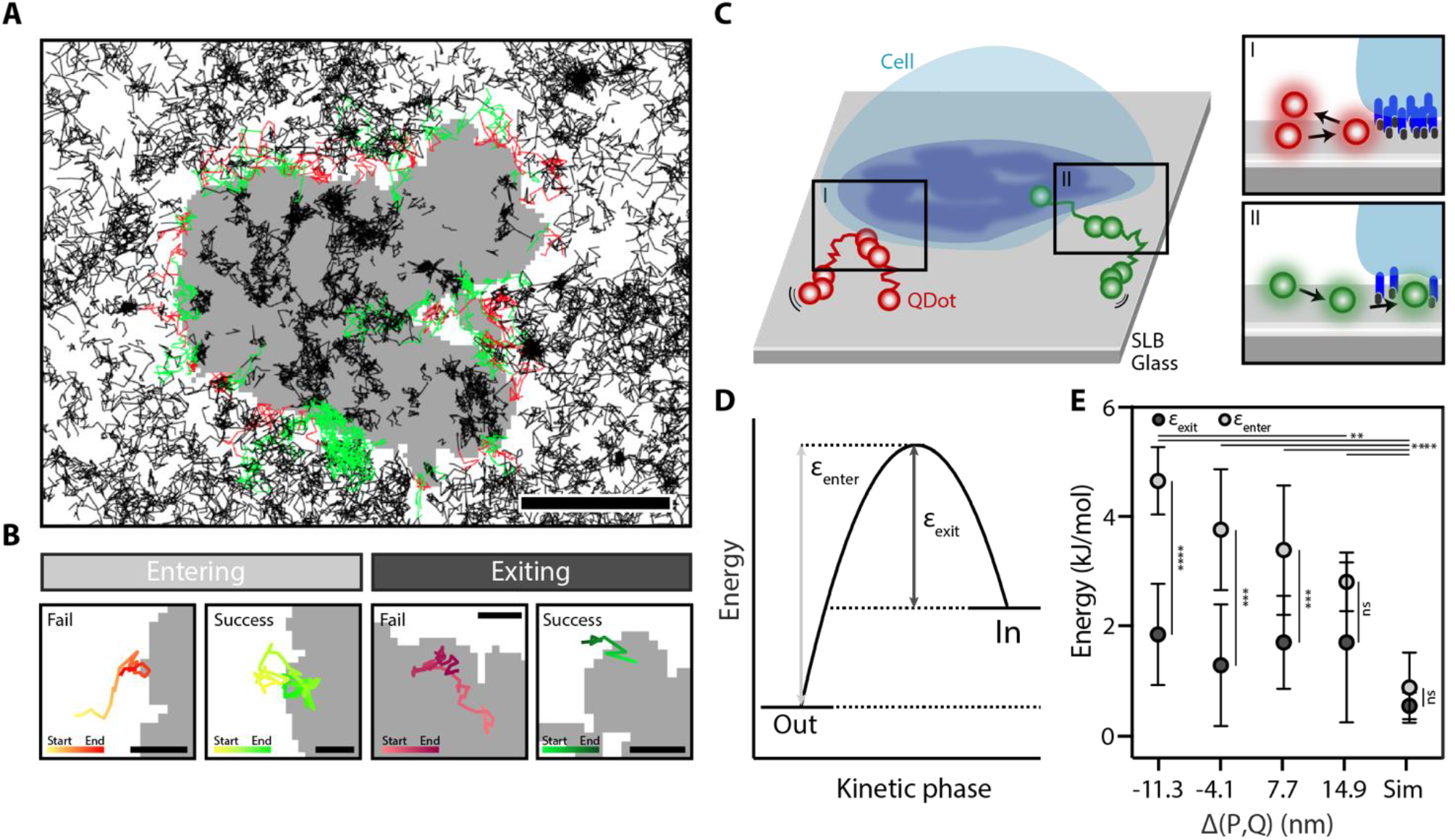
Energy penalty of contact entry/exit for QDots. (A) Tracks are marked depending on their success in entering the contact (Black: tracks not attempting to enter or exit the contact, green: success in entering the contact, red: fail to enter the contact). (Scale bar 5 µm). (B) Examples of tracks failing and succeeding to enter (left) or exit (right) the contact. (Scale bar 1 µm) (C) Schematic illustration of QDots attempting to enter the cell contact ((I) fail: red, (II) success: green,). (D) Energy landscape for entering/exiting the contact. (E) Energy penalty for entering (ε_enter_) and exiting (ε_exit_) for different gap size conditions calculated via success and failed attempts assuming a Boltzmann distribution (mean±s.d.). P values were obtained from a one-way ANOVA test and are shown for the success rate in entering. *p < 0.1, **p < 0.01, ****p < 0.0001. The p values above the graphs are energy penalties for entering QDots and below for exiting QDots. Sim shows results for a simulation with 20 repeats and two different contacts. In the simulation, energy penalties are calculated for tracks starting outside and entering the contact (ε_enter_) and tracks starting inside the contact and exiting (ε_exit_). Number of cells n for each condition: Δ(P,Q)=−11.3 nm, n=10, Δ(P,Q)=−4.1 nm, n=6, Δ(P,Q) =7.7 nm, n=10, Δ(P,Q) =14.9 nm, n=7. Find all number of attempts, success and failures in Suppl. Table 1.

To further verify the increased entering energy penalty for QDots larger than the contact size, we compared ε_enter_ for the measured contact barrier with an area surrounding the contact (dilated mask) in the bilayer (see Suppl. Fig 4), where we would not expect entry restrictions. The result confirmed that the observed energy penalty is due to the contact edges acting as a barrier, since there is no entry restriction in the dilated masks compared to the contact (see Suppl. Fig 4).

### QDot slowdown in contacts is independent of size

After having established the contact edge as a barrier we now focus on the heterogeneity of the contact. Here, areas of reduced velocity were located by studying the behaviour of QDots inside the cell contact, measuring the instantaneous velocity or jump-distance (JD) of the probes between frames. The steric hindrance landscape was visualised using a map of the average JD of QDots for each pixel (Fig. 3A). We observed a reduction in JD for tracks inside the cell contact, implying steric hindrance, restricted movement or reduced mobility in the contact. By fitting the JD distribution, we found diffusion coefficients D between 0.49±0.05 µm^2^/s and 0.83±0.03 µm^2^/s (mean±s.d.) for QDots diffusing outside the cell contact (Fig. 3B). These values are in agreement with reported lipid diffusion in bilayers, which ranges from 1 to 4 μm^2^s^−1^ (29). The mean velocity of the QDots inside the contact was reduced ~2 fold versus that for diffusing QDots outside the contact (Fig. 3B). Interestingly, the mean velocity inside the contact was constant over all gap-size conditions assessed (Fig. 3B), although we saw a slight shift towards lower velocities in the JD distributions for negative Δ(P,Q) (Suppl. Fig. 5).

**Figure 3.**
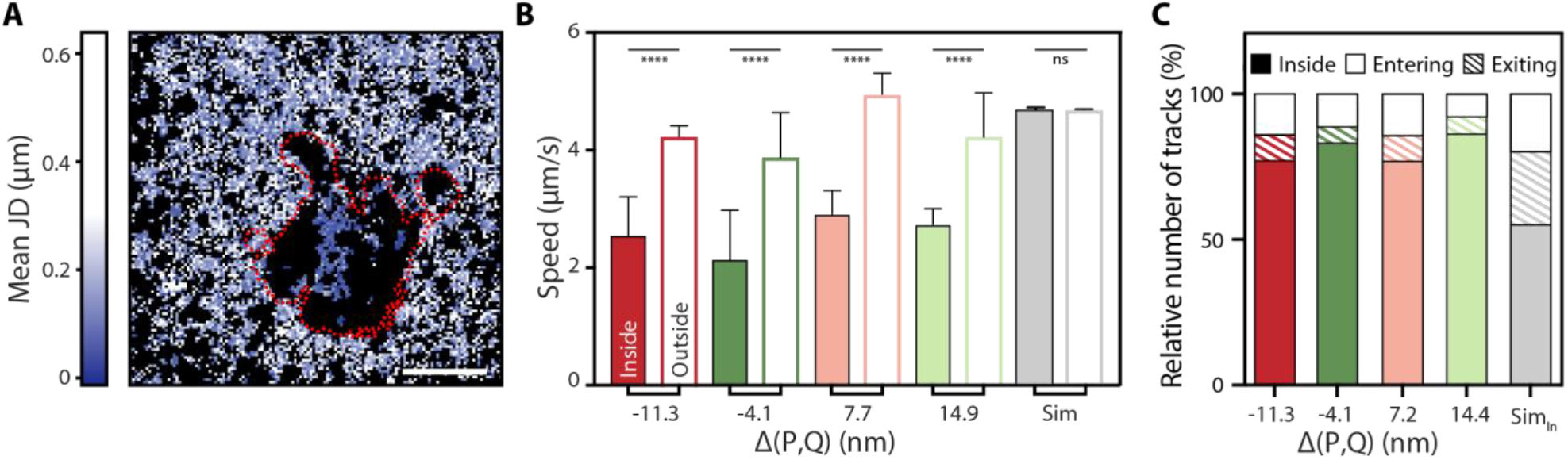
Gap-size dependent slow-down of QDots. (A) Jump distance (JD) map of a cell-bilayer contact. Image shows a representative cell of CD2CD45 spacer proteins used with Q605 diffusing in the SLB. The red dashed line represents the outline of the cell contact. Scale bar, 5 µm. (B) Average speed of QDots inside and outside contact in µm/s. Error bars represent mean±s.d. Sim shows results for a simulation with 20 repeats and two different contact masks. Tracks were either starting inside (Sim_in_) or outside the contact. Number of cells n per experiment: Δ(P,Q)=−11.3 nm, n=10, Δ(P,Q)=−4.1, n=6, Δ(P,Q)=7.7 nm, n=10, Δ(P,Q) =14.9 nm, n=7. ****p < 0.0001 (C) Number of tracks entering (open) and exiting (shaded) the cell gap in comparison with tracks, which are already inside the cell contact (filled).

Overall, we observed a trapping effect for all probes, that was gap-size independent. We found that 80% of QDots do not leave the contact over the time of acquisition, which is significantly different from simulations (50% remain) (Fig. 3C). This difference between simulation and experiment reflects the larger energy penalty ε_exit_ in experimental data compared to the simulations, which can be explained by the complete absence of protein crowding effects in the simulation data. In experimental data, probes in the contact might experience crowding, hindering their exit, gap-size independently. Assuming similar restrictions hold for proteins of comparable size, the contact is a very exclusive environment with exchange limited to ~20%. If we extrapolate these to signalling molecules, this size-independent trapping could lead to a prolonged residence time inside the contact, exposing them, in the case of T-cell synapses, to increased kinase to phosphatase ratios.

Although we did not see an overall effect of gap size on QDot velocity, the data clearly revealed a reduction of mean velocity inside compared to outside the contact (Fig. 3B). Interestingly, the reduction in mean velocity was independent of the protein spacer size, suggesting that the molecules were hindered in the contact due to effects such as crowding. The observed differences in energy penalties are not a result of differences in contact areas, as all contact areas were comparable (see Suppl. Fig. 6.). In addition, we observed areas in the contact that seemed inaccessible to the probes (see Fig. 3B). We observed spacer accumulation, implying the formation of a local gap of known height, with no QDot tracks underneath. Here, the probes were mostly deflected out of the contact, with emerging “pathways” along which probes diffuse once inside the contact. These pathways could have been indicative of membrane ruffles below the diffraction limit.

To determine if some areas of the contact border were easier to access than others, we identified “breach zones” i.e. parts of the boundary where QDots could successfully cross the border (Suppl. Fig. 7). Here, we found a predominantly negative correlation between entry success rate and fluorescence intensity of the spacer protein for all gap size conditions (Suppl. Fig 7). In other words, we find lower success rates in areas of higher contact intensity, implying that the protein spacers create a physical barrier to entry simply due to the increased density of proteins in the contact versus outside. This effect of protein spacer density on the probe prompted us to study inhomogeneities of spacer protein density and the effect on probe diffusion.

### QDot diffusion coefficient is reduced in a contact density-dependent manner

Since the studied cell-bilayer contacts were not homogenous, but the distribution of the spacer protein varied across the contact, not only from cell to cell due to expression level variation, but also within each contact for a given cell, this allowed us to compare regions of different spacer protein density within one cell. To investigate if hindered diffusion in the contact scales with spacer protein density, we studied the correlation between pixel intensity for the protein spacers and the average jump distance of the QDot measured for a given pixel (Fig. 4A). One would expect a higher intensity to correspond with a higher density of spacer protein, implying a more stable or uniform cell/bilayer contact. In these regions QDots should experience greater steric hindrance. To derive this correlation, we divided the contact area into regions according to their spacer protein pixel intensities and overlaid these with the JD maps (Fig. 4B, see JD curves in Suppl. Figure 8). Fitting the resulting JD distributions (22), we found that the highest protein spacer density corresponded to the lowest diffusion coefficient, for all gap size conditions examined (Fig. 4 C, D). We observed an almost linear decrease in diffusion coefficient with protein density for negative Δ(P,Q). For the larger CD2CD45-mediated gap the reduction in QDot diffusion was not as pronounced (Fig. 4D). However, this effect was more pronounced when comparing the behaviour of the same QDot in different gap heights. Here, the diffusion of the small QDot (Q525) was reduced by 85% when compared with QDots outside the contact for areas of negative Δ(P,Q), whereas it was only halved for regions of positive Δ(P,Q). This suggests, that both the gap-height and the density of adhesion proteins are involved in shaping the behaviour at the contact.

**Figure 4.**
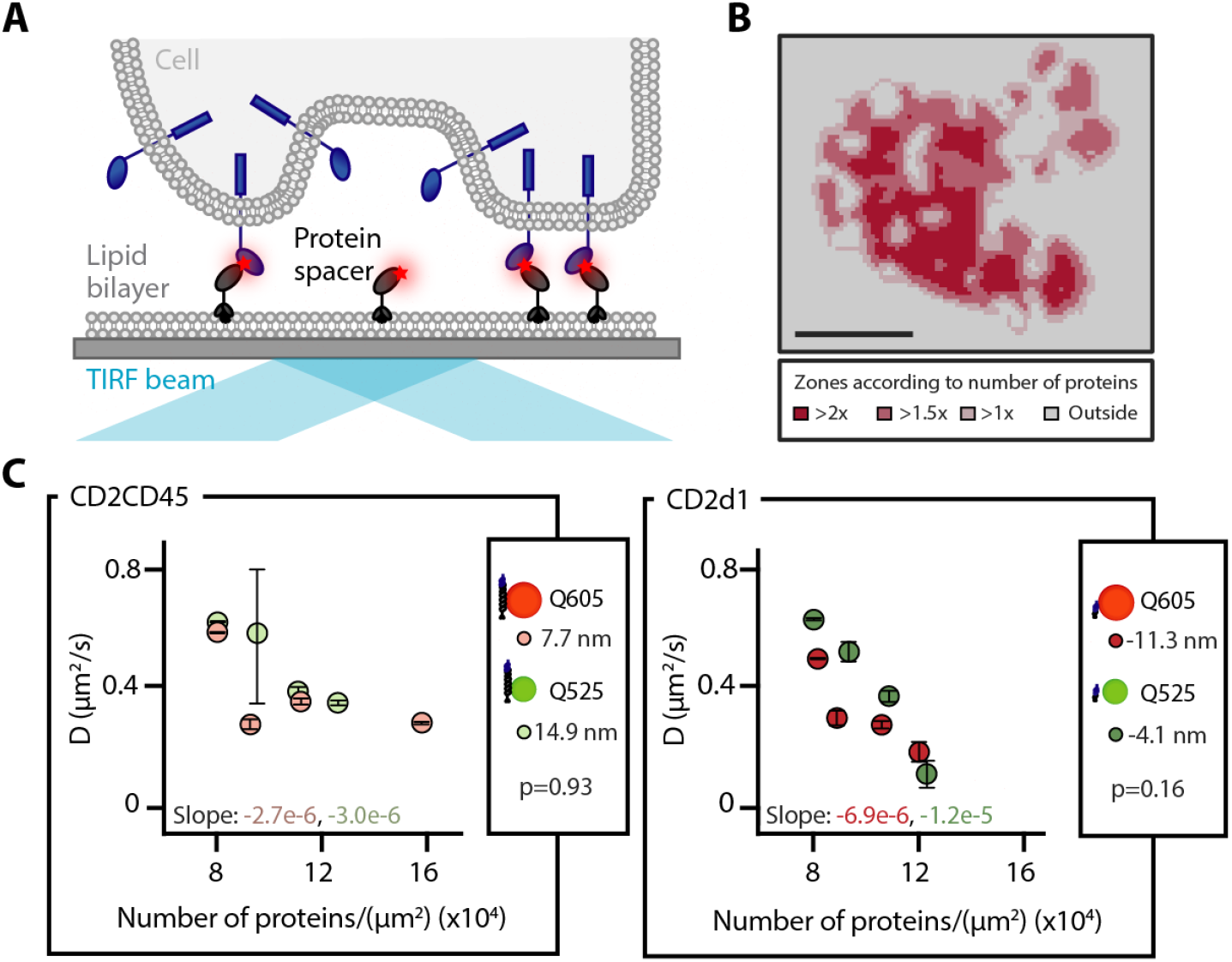
QDot Diffusion influenced by spacer density. (A) Schematic representation of protein spacers within the contact. (B) Intensity zones of the labelled protein spacer in different areas of the contact. Scale bar, 5 µm. (C) Diffusion constants for QDots in areas of different protein spacer density, comparing diffusion coefficient (D) values for constant spacer and QDot size. Slope values were analysed using a one-way ANOVA (see Suppl. Fig. 9). Values represent the mean, error bars were obtained with boot strapping (number of iterations = 1000). Δ(P,Q)=−11.3 nm, n=10, Δ(P,Q)=−4.1 nm, n=6, Δ(P,Q) =7.7 nm, n=10, Δ(P,Q) =14.9 nm, n=7.

While the protein spacer density could increase steric hindrance due to crowding, it could also promote more homogenous gap height, as the spacer proteins we used were expected to be relatively rigid. Both effects, crowding and gap-size could influence diffusion and access into the contact. Since the bilayer intensities between all conditions were similar (see Suppl. Fig. 1), we could compare between the conditions, assuming saturation of the nickelated groups. We found that diffusion was decreased for larger QDots in smaller gaps, while smaller probes in larger gaps seemed to be less affected by protein density (Fig. 4). This behaviour most likely also applies to proteins in the T-cell contact, as most T cell membrane proteins are heavily glycosylated rendering their conformation stiff and extended (30). Interestingly, Douglass and Vale identified signalling islands on T cells where important signalling proteins are reduced in their diffusion through protein protein interactions, forming so called microdomains (13). Here, the proteins studied did not possess an extracellular domain which could impose steric hindrances, but a slow-down of up to 50% could still be observed. In our study, we found that even without any binding effects of the probe similar trapping effects can be observed through increased adhesion protein density. While we studied the impact of steric effects on the extracellular domains of proteins, a similar effect of intracellular crowding due to increased protein density cannot be excluded and is of interest for future studies.

## Conclusion

We present a method to study inhomogeneities in cellular contacts, demonstrated by following QDots in contacts formed by T cells interacting with a supported lipid bilayer, at single-particle densities. We measured the density and instantaneous velocity of QDots inside and outside contacts, determined the energy penalty to enter and exit the contact and created spatial maps of the diffusion rates and jump distances which could then be compared to the density of the protein spacers. The results provided a detailed view of the cell-bilayer contact and suggested that significant levels of trapping of large molecules could occur, with relatively little exchange of molecules across the contact boundary and a significant decrease in diffusion, once the contact forms. We found a gap-size dependent energy penalty for entering contacts and a gap-size independent penalty for exiting. Furthermore, the contact and boundary are not uniformly accessible, with accessibility scaling with spacer protein density. We also observed a protein spacer density-dependent slowing down of QDots across all gap sizes. This approach has the potential to reveal the likely behaviour of molecules at cell-contacts, elucidating how and why their diffusion and exchange with molecules outside the contact is modified, and how such effects could influence signalling.

## Supporting Material

Svideo 1A and 1B. QDots 605 diffusing in a bilayer and through a CD2rCD45 cell contact. (A) Raw video of QDots diffusing in bilayer. (B) Time-lapse of tracks colour-coded according to track speed (analysed with TrackMate by Fiji)

Svideo 2. QDots 525 diffusing in a bilayer and through a CD2rCD45 cell contact

Svideo 3A and 3B. Simulation of QDots diffusing freely in a bilayer and through a CD45 cell contact. (A) Tracks starting inside the cell contact. (B) Tracks starting outside the cell contact

Suppl. Table 1: Number of tracks attempting to enter or exit the cell contact and success to enter or exit.

Suppl. Fig 1: Variation of bilayer intensity for different conditions

Suppl. Fig 2: QDots diffuse freely in bilayer (MSD plot)

Suppl. Fig 3: Shift of JD distribution between attempts, exit and enter

Suppl. Fig 4: Energy penalties for eroded and dilated contact borders

Suppl. Fig. 5: ECDFs of overall jump distances in track image inside the contacts

Suppl. Fig. 6: Contact sizes between gap conditions similar

Suppl. Fig 7: Negative correlation between success rate and spacer pixel intensity

Suppl. Fig 8: JD distributions separated out via spacer intensity zones

Suppl. Fig 9: Additional Statistics for linear fit of diffusion constants in areas of various protein density.

## Author contributions

A.H.L., K.K. and D.K. designed the experimental plan. A.M.S. provided cell lines and purified proteins. K.K. and A.H.L. performed experiments and data analysis. A.P. provided the tracking simulation code. A.H.L., K..K., J.M., A.M.S., A.P., S.F.L., S.J.D and D.K. wrote the article.

## Acknowledgements

This work was supported by a Royal Society University Research Fellowship (UF120277 to S.F.L.) and a Research Professorship (RP150066 to D.K.); the EPSRC (EP/L027631/1 to A.P.); and the Wellcome Trust (098274/Z/12/Z to S.J.D.).

